# Delivery of Circular mRNA via Degradable Lipid Nanoparticles against SARS-CoV-2 Delta Variant

**DOI:** 10.1101/2022.05.12.491597

**Authors:** Ke Huang, Na Li, Yingwen Li, Jiafeng Zhu, Qianyi Fan, Jiali Yang, Yinjia Gao, Yuping Liu, Qiangbo Hou, Shufeng Gao, Ke Wei, Chao Deng, Chijian Zuo, Zhenhua Sun

## Abstract

mRNA vaccines have emerged as a most promising and potent platform in the fight against various diseases including the COVID-19 pandemic. However, the intrinsic instability, varying side effects associated with the delivery systems, and continuous emergence of virus variants highlight the urgent need for the development of stable, safe and efficacious mRNA vaccines. In this study, by screening a panel of proprietary biodegradable ionizable lipidoids, we reported on a novel mRNA vaccine (cmRNA-1130) formed from a biodegradable lipidoid with eight ester bonds in the branched tail (AX4) and synthetic circular mRNA (cmRNA) encoding the trimeric Delta receptor binding domain (RBD) of SARS-CoV-2 spike protein for the induction of robust immune activation. The AX4-based lipid nanoparticles (AX4-LNP) revealed much faster elimination rate from liver and spleen in comparison with commercialized MC3-based LNP (MC3-LNP) and afforded normal level of alanine transferase (ALT), aspartate aminotransferase (AST), and creatinine (CRE) in BALB/c mice. Following intramuscular (IM) administration in BALB/c mice, cmRNA-1130 elicited potent and sustained neutralizing antibodies, RBD-specific CD4^+^ and CD8^+^ T effector memory cells (Tem), and Th1-biased T cell activations. cmRNA-1130 vaccine showed excellent stability against 6-month storage at 4 °C and freezing-thawing cycles. In brief, our study highlights mRNA vaccines based on cmRNA and biodegradable AX4 lipids hold great potential as superb therapeutic platforms for the treatment of varying diseases.

## INTRODUCTION

Coronavirus disease 2019 (COVID-19) is a serious global public health emergency caused by the novel severe acute respiratory syndrome coronavirus (SARS-CoV-2)^[1]^. According to report from the World Health Organization (WHO), COVID-19 has so far caused nearly 500 million confirmed cases and more than 60 million confirmed deaths. On top of that, SARS-CoV-2 continues evolving into many variants, and many of these variants with evidence to enhance viral transmissibility, infectivity, and to escape from host immune response are classified as variants of concerns such as Delta variant of SARS-CoV-2 (also known as B.1.617.2)^[2]^. The Delta variant is highly transmissible, reported to be more than twice as transmissible as the original strain of SARS-CoV-2^[3]^. Therefore, there is an urgent need to develop safe and effective vaccines against SARS-CoV-2 Delta infection. Variety categories of vaccine programs have been initiated using a variety of SARS-CoV-2 antigen and vaccine modalities, inactivated/live attenuated virus, recombinant adenovirus, recombinant protein, DNA vaccine, and mRNA vaccine^[4]^. It is worth noting that the mRNA vaccine has demonstrated protective capacity and promising immunogenicity in animal and humans^[5]^. The mRNA platform offers several benefits over other nucleic acid-based therapeutics as mRNA is non-infectious, does not require transport to the nucleus and integrate into the host genome.

Compared to the linear mRNA, circular RNA (circRNA) is highly stable due to its covalently closed ring structure, which protects it from exonuclease-mediated degradation^[5a, 6]^. It has been reported that circRNAs were more stable than their linear mRNA counterparts, with the median half-life was at least 2.5 times longer than their linear mRNA isoforms in mammalian cells^[6-7]^. Recent studies revealed that circRNAs could be used as vectors for protein expression^[8]^. In previous studies, we developed a novel circular mRNA platform, termed “cmRNA”, to mediated potent and durable protein expression *in vitro* and *in vivo*^[7c]^. We proved that cmRNA directed higher and more durable protein expression than the typical linear mRNA, and could be applied to express different proteins and cytokines for cancer immunotherapy. The *in vivo* efficacy of cmRNA as therapeutic agent or vaccine can be boosted by the introduction of efficient and safe delivery systems.

In recent years, lipid-based nanovehicles like lipid nanoparticles (LNP), have shown significant potential in the delivery of mRNA, with high safety and effectiveness in clinical trials^[7a, 9]^. In 2018, the U.S. Food and Drug Administration (FDA) approved Onpattro, a LNP delivered small interference RNA (siRNA) drug targeting transthyretin, for the treatment of amyloidosis^[10]^. Two year later, mRNA-LNP based SARS-CoV-2 vaccines from Moderna and Pfizer/BioNTech received emergency use authorization (EUA) in several markets^[9b, 9c, 10]^. However, it has been reported that the developed LNP might induce highly inflammatory, causing obvious side effects or potential toxicities^[11]^. Therefore, novel LNP delivery platform with better biodegradability and biocompatibility are urgently required.

Here, we developed a new SARS-Cov-2 Delta variant vaccine, named cmRNA-1130, in which the cmRNA encoding Delta RBD trimer antigen was encapsulated by a novel biodegradable LNP. The LNP is composed of a leading tail-branched, eight ester bonds included biodegradable lipidoid (AX4) and three coformulated excipient lipid molecules. The successful *in vivo* delivery of delta RBD cmRNA elicited robust neutralizing antibodies (NAbs) and strong T cell responses in mice. Moreover, NAbs elicited by cmRNA-1130 vaccination exhibited notable neutralizing activities against Delta variant of SARS-CoV-2 pseudovirus, and the NAbs in mice can maintain at least 132 days, suggesting that the protective effect may be highly efficient and durable. Additionally, this vaccine can be stored at 4°C for more than 6 months. Collectively, our findings manifesting the huge clinical potential of cmRNA-1130 as a prophylactic vaccine to directly blockade COVID-19 pandemic.

## RESULTS

Ionizable lipid materials with low toxicity and facile synthesis have attracted enormous attention for the construction of mRNA delivery systems. The incorporation of degradable linkers like ester bond into the aliphatic tails of lipidoids often diminishes the immunogenicity, facilitates mRNA release, and improves mRNA expression^[12]^. Furthermore, tail-branched lipids have been reported to accelerate their escape from endosomes^[12c, 13]^. Here, a series of ionizable lipids with multiple ester bonds in the branched tails were readily synthesized by Michael addition reaction (**Figures 1A & 1B**). The structure of a typical lipid AX4 was verified by ^1^H NMR spectra (**Figure S1**). To evaluate the vaccination potential, these ionizable lipids were formulated into LNP along with the excipient components, cholesterol, 1,2-distearoyl-sn-glycero-3-phosphocholine (DSPC), and 1,2-dimyristoyl glycerol-*rac*-3-methoxy-poly(ethylene glycol)-2000 (DMG-PEG), and maintained same molar composition and nitrogen-to-phosphate ratio (N:P). mRNA delivery efficacy was evaluated by loading firefly luciferase (Fluc) circular mRNA as a reporter into LNP, and then comparing luciferase protein expression in mice following intramuscularly (IM) administration. Mice treated with Fluc mRNA-LNP revealed that ionizable lipids with a tail containing butyloctanoic acid (X4) had better mRNA delivery and higher protein expression than lipids with hexyldecanoic acid (X6) (**Figure 1C**). As for the head of lipidoids, tertiary amine with two amines in the polar head group (A) facilitated the introduction of four branched tails at most and demonstrated superior delivery efficacy, in which AX4 and AX6 showed 2- and 0.2-fold higher protein expression than commercialized Dlin-MC3-DMA lipid (MC3), respectively. Of note, mice administrated with AX4 exhibited sustained luciferase expression within 24 h, and the luciferase reached the maximum at 6 h post-injection (**Figures 1D & S2**). The inclusion of heterocyclic structures (B & C) induced moderate delivery efficiency except that lipid (DX4) with imidazolyl D afforded 1-fold higher protein expression in comparison with MC3. Incorporating ester, ether, tertiary amine, and hydroxyl to the lipid head afforded less protein expression than MC3, while the extended space between hydroxyl and amine appeared to enhance delivery efficacy. Based on these preliminary results, AX4 was recognized as the most potent mRNA delivery material.

**Figure 1.**
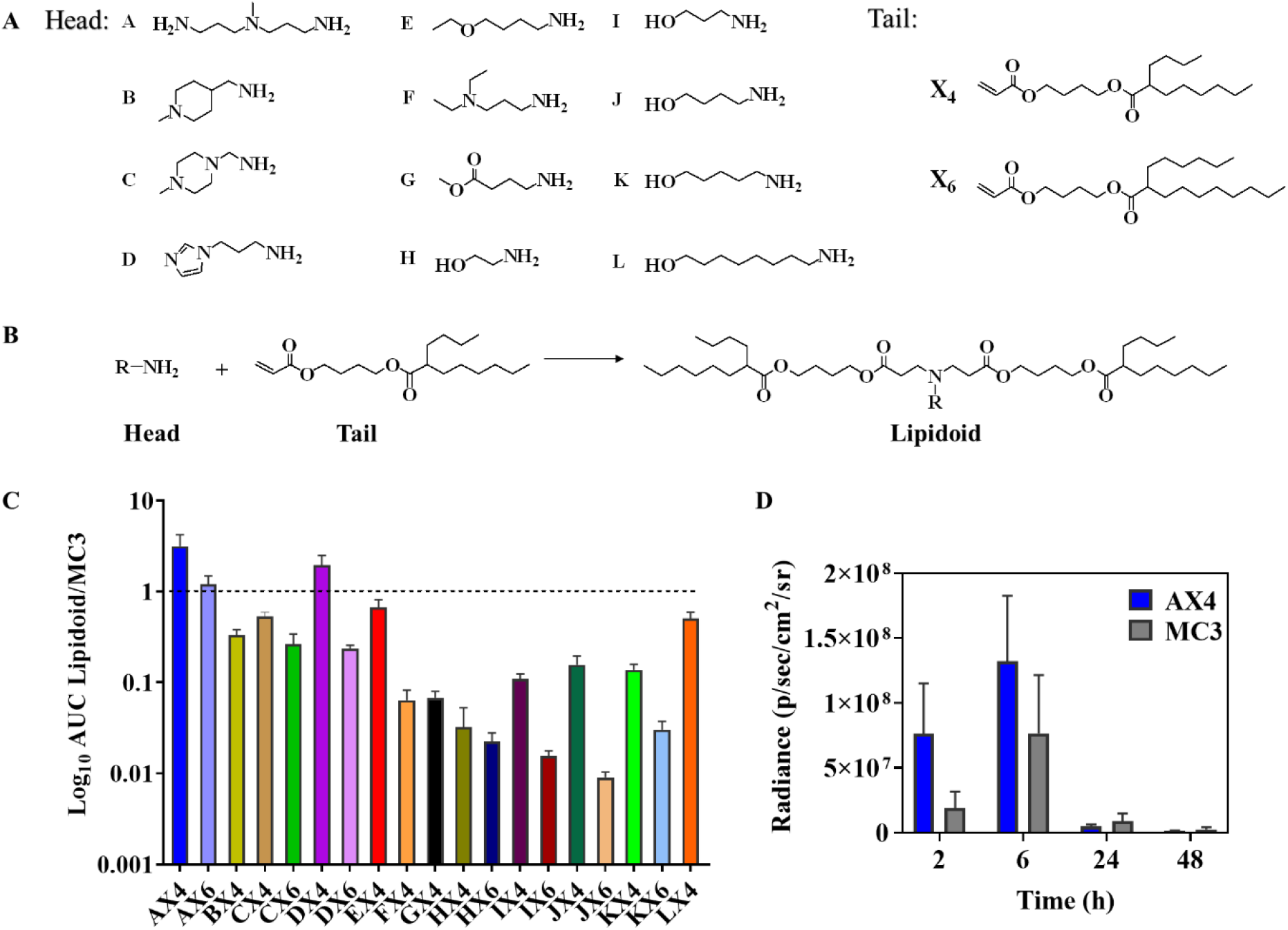
*In vivo* screening of lipidoids for mRNA delivery. (A) Chemical structures of head and tail used for lipidoid synthesis. (B) Synthesis of lipodiods through Michael addition reaction between amine and acrylate. (C) Area under curve (AUC) assay of whole-body luciferase bioluminescence of BABL/c mice following IM injection with AX4-LNP at a dose of 0.5 mg Fluc mRNA equiv./kg for 2 h (n = 3). (D) Quantification of total luciferase expression in mice IM administered with AX4-LNP or MC3-LNP for 2, 6, 24 and 48 h (n = 3).

To investigate the effect of tail numbers of lipidoids on mRNA expression, AX4-2 with two branched X4 and AX4-3 with three X4 were designed and synthesized (**Figure 2A**). The results showed that increasing the number of branched X4 afforded enhanced protein expression levels, in which AX4 exhibited three times and three orders of magnitude higher protein levels than AX4-3 and AX4-2, respectively (**Figure 2B**). The significant declined protein expression of AX4-2 could attribute to that increased secondary amines would hamper mRNA release inside cells through the formation of relatively strong electrostatic interactions.

**Figure 2.**
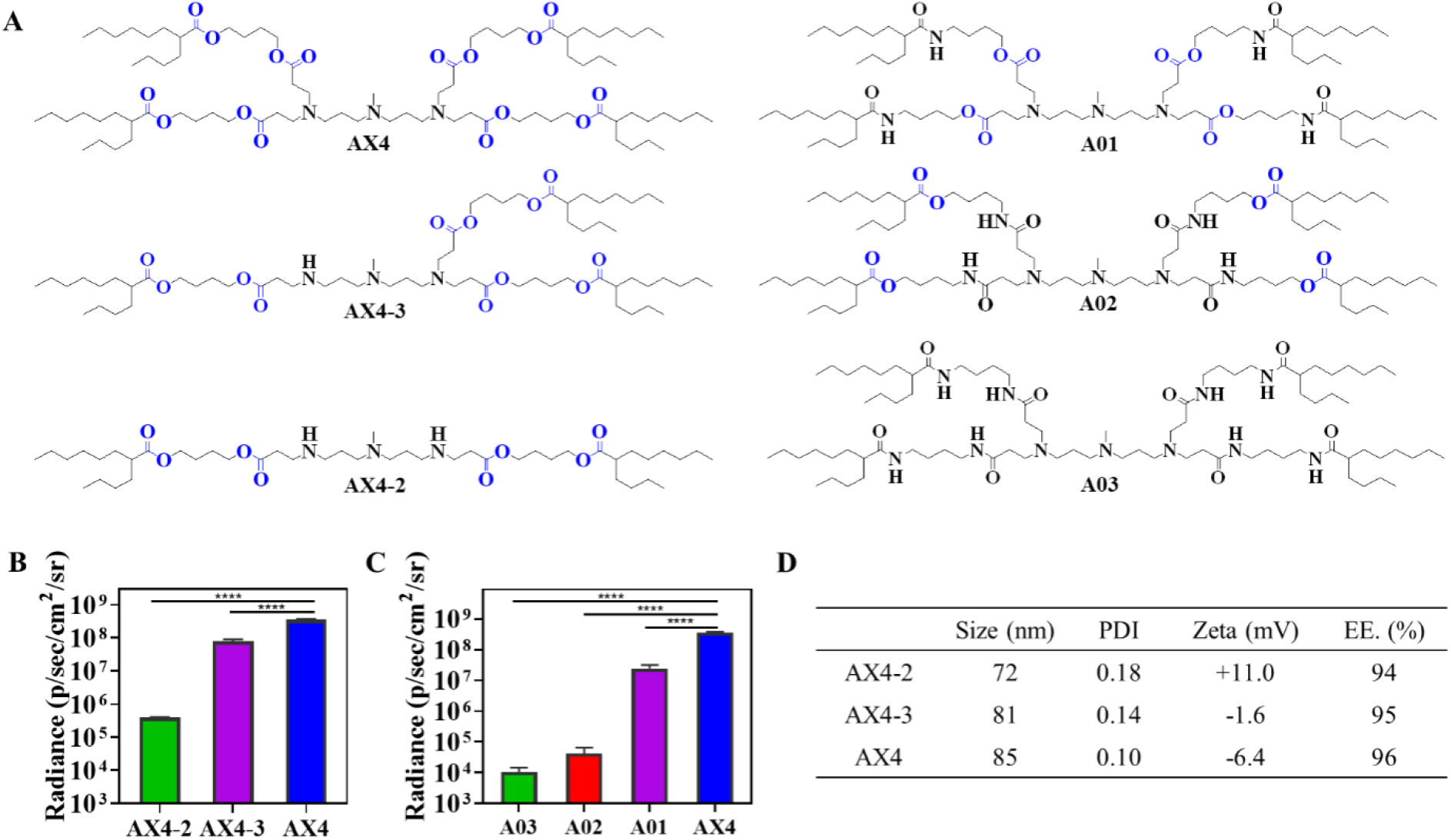
*In vivo* screening of lipidoids with different linkers and varying number of X4 for mRNA delivery. (A) Structures of ionizable lipidoids. (B-C) Quantification of total luciferase expression in BABL/c mice following IM administration of Fluc mRNA-LNP at a dose of 0.5 mg mRNA equiv./kg for 6 h (n = 3). \**P* < 0.05, ** *P* < 0.01, *** *P* < 0.001, **** *P* < 0.0001, n.s. = not statistically significant. (D) Physicochemical properties of Fluc mRNA-LNP, including the size, PDI, zeta potential, encapsulation efficiency (EE.).

To understand the role of the ester bond in the lipidoid tails, more stable amide bonds were employed to replace part (A01, A02) or all (A03) of the ester bonds in AX4, as shown in **Figure 2A**. The incorporation of amide bonds in lipids appeared to significantly decrease the luciferase intensity, mainly owing to that the amide bonds retarded the lipid degradation and encumbered the mRNA release (**Figure 2C**). Moreover, the replacement of amide bonds away from tertiary amide (A01) displayed much more protein expression than that adjacent to tertiary amide (A02), partly owing to that A01 following the degradation of ester bonds provided shorter alkyl chains and carboxyl groups to facilitate mRNA release. The increase of amide bonds provided lipids with less degradability, and lipids with eight amide bonds (A03) revealed lowest luciferase expression level. Fluc mRNA-encapsulated AX4-LNP (Fluc mRNA-AX4-LNP) displayed a mean diameter of about 85 nm, narrow distribution with a polydispersity index (PDI) of 0.10, and slightly negative charge (−6.4 mV). Decreasing the number of X4 in the lipidoids afforded smaller sizes and higher surface charges (−1.6 ~ 11.0 mV). Of note, over 94% encapsulation efficiency (EE.) of mRNA was achieved for all LNP formed from AX4, AX4-2, and AX4-3.

mRNA labeled with Cy5 (Cy5-mRNA) was employed to determine the *in vivo* biodistribution of mRNA-AX4-LNP. Strong Cy5 fluorescence was detected in the liver, kidney, and gallbladder at 6 h post-injection, regardless of intravenous (IV) and IM administration (**Figure 3A**). However, significant luciferase fluorescence was only observed in the whole liver (**Figures 3A-C**), signifying the liver-targeted protein expression of mRNA-AX4-LNP. It is well known that the liver-targeting LNP acquired of ApoE in the serum results in uptake of lipoproteins by liver hepatocytes through receptor-mediated endocytosis by low-density lipoprotein receptor (LDL-R)^[10, 14]^. To explore whether the endogenous ApoE affects the protein expression level of mRNA-AX4-LNP in different organs, a ApoE knockout (ApoE^−/−^) mouse model was employed. In sharp contrast with the brilliant luciferase fluorescence in wild-type C57BL/6 mice, mRNA-AX4-LNP IV administration induced little protein expression in ApoE^−/−^ mice with around 90% reduction of luciferase intensity (**Figures 3D & E**).

**Figure 3.**
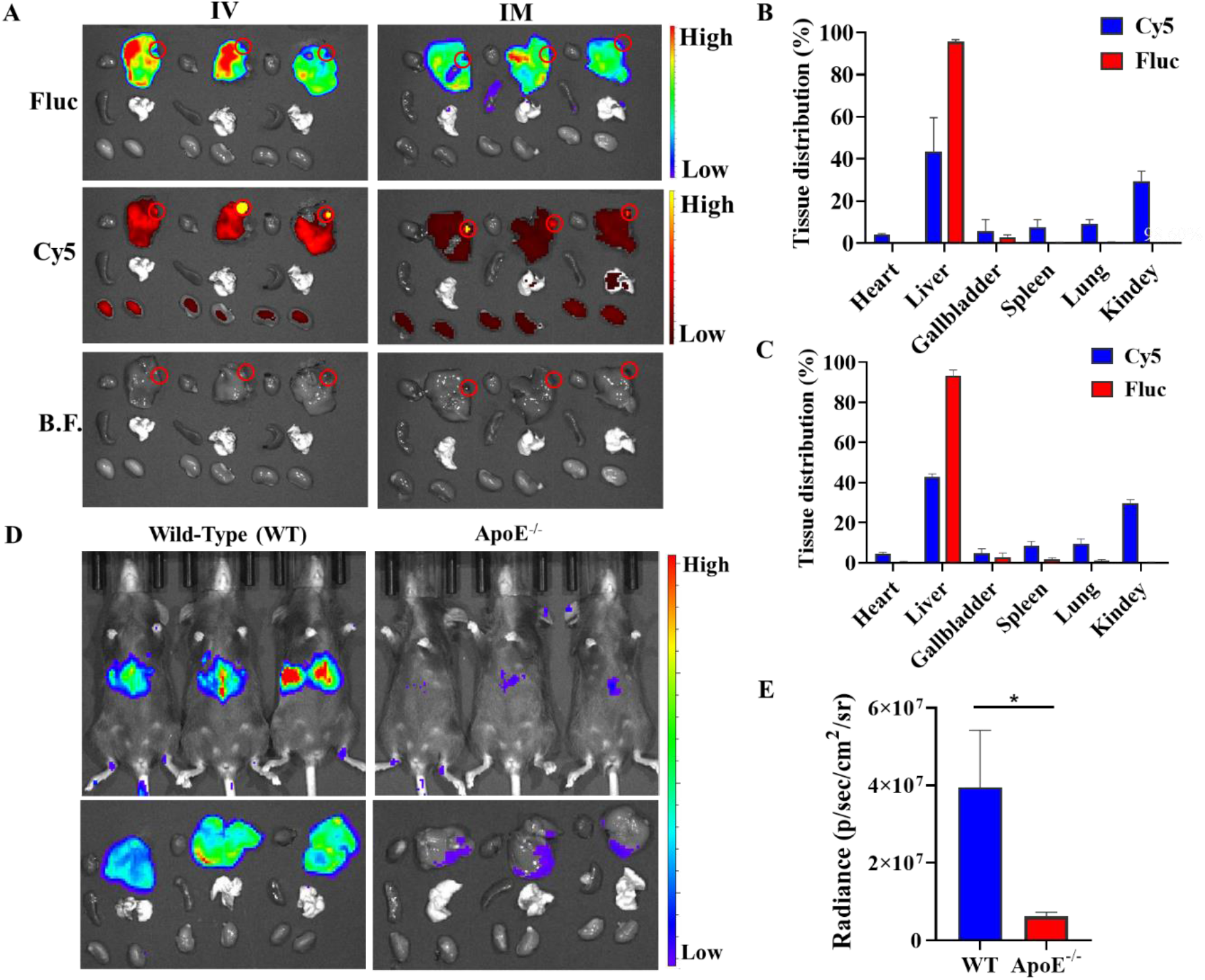
*In vivo* biodistribution and protein expression of Fluc mRNA-AX4-LNP following I.M or intravenous (IV) administration. (A) Fluorescence in major organs detected at 6 h after IM or IV injection of Fluc mRNA-AX4-LNP (n = 3). Cy5 and Fluc represented mRNA biodistribution and expression in main organs, respectively. The red circles indicated the gallbladder sites. (B-C) Quantitative analysis of *in vivo* biodistribution and protein expression following IV (B) and IM (C) administration. (D) Comparison of protein expression in wild type (WT) mice and ApoE^−/−^ mice at 6 h following IV injection of Fluc mRNA-AX4-LNP (n = 3).(E) Quantification of total luciferase expression. \**P* < 0.05, \*\**P* < 0.01, \*\*\**P* < 0.001, \*\*\*\**P* < 0.0001, n.s. = not statistically significant.

AX4 was formulated into LNP by mixing with DSPC, cholesterol and DMG-PEG, and Fluc mRNA via microfluidic technique. The formed mRNA-AX4-LNP revealed an average particle diameter of 85 nm (**Figures 4A & 4B**). The Cryo-electron microscopy images highlighted the relatively uniform spherical morphology of AX4-LNP (**Figure 4B**). As shown in **Figure 4C**, AX4-LNP revealed an acid dissociation constant (pKa) of 6.89, facilitating the endosomal escape and mRNA release of the formed LNP in targeted cells.

**Figure 4.**
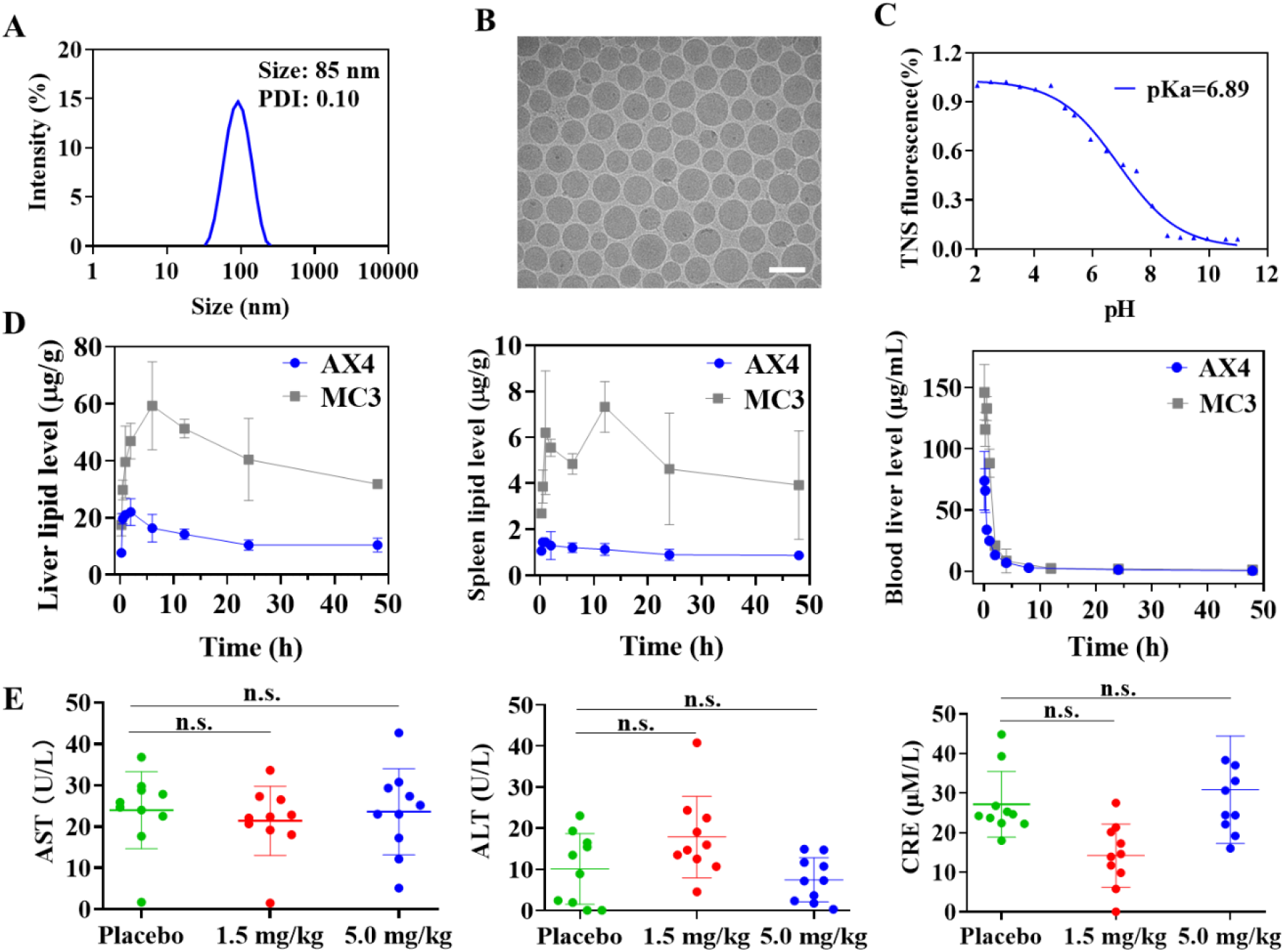
(A) Intensity size of mRNA-AX4-LNP measured by dynamic light scattering. (B) Cryo-electron microscopy image of mRNA-AX4-LNP. Scale bar, 100 nm. (C) pKa of AX4-LNP determined by TNS method (n = 3). (D) Metabolic behaviors of AX4-LNP and MC3-LNP with time in liver, spleen and blood following IV administration (n = 3). (E) Serum level of alanine transferase (ALT), aspartate aminotransferase (AST), and creatinine (CRE) in BALB/c mice that received AX4-LNP coformulated with RBD mRNA at two doses of 1.5 and 5.0 mg/kg, PBS was used as a control. Serum was collected at day 14 after secondary injection (n = 10). \**P* < 0.05, \*\**P* < 0.01, \*\*\**P* < 0.001, \*\*\*\**P* < 0.0001, n.s. = not statistically significant.

The metabolic behaviors of AX4-LNP in liver and spleen was monitored by LC-MS/MS. In comparison with MC3-LNP, AX4-LNP revealed a much faster elimination speed from liver and spleen following IV administration (**Figure 4D**). For example, MC3 revealed a maximum concentration (C_max_) of 59 μg/g in the liver at 6 h post-administration, and decreased to 33 μg/g when the time extended to 48 h. In comparison, AX4 displayed a small amount of residual in the liver during the whole experiment process with a C_max_ of 21 μg/g at 2 h post-administration. Much less residual in the spleen was observed for both AX4 and MC3 comparing with that in the liver. In addition, AX4 showed less than 10 μg/g residual within 48 h, which was around three times lower than MC3. The fast metabolic behaviors of AX4 would decrease the potential systemic toxicity associated with delivery systems and facilitate possible repeatable dosing of mRNA vaccines. MC3 and AX4 revealed similar elimination kinetics and could be removed from the blood within 8 h.

Intravenous (IV) injection of AX4-LNP in BALB/c mice at doses of 1.5 and 5.0 mg mRNA equiv./kg revealed similar level of alanine transferase (ALT), aspartate aminotransferase (AST) and creatinine (CRE) in serum comparing with PBS group, verified that mRNA vaccine caused negligible impairment to liver and kidney (**Figure 4C**). In addition, mice inoculated with mRNA vaccine demonstrated no obvious body weight change (**Figure S3**).

According to the previous studies, the LNP formulated SARS-Cov-2 RBD mRNA vaccine (BNT162b1) elicited high level of spike/RBD neutralizing antibodies and potent strong T-helper-1 CD4^+^ and IFNγ^+^ CD8^+^ T cell responses in pre-clinical studies, and ultimately resulted in successful protection of rhesus macaques from the challenge of SARS-Cov-2 ^[15]^. Later studies of this vaccine candidate in Phase I clinical study revealed good tolerance in adults, and high level of neutralizing antibodies, as well as IFNγ^+^ CD8^+^ T cell responses were confirmed^[16]^. These results suggest that RBD trimer is sufficient for SARS-Cov-2 vaccine. In this study, RBD (residues Arg319–Phe541) of wild-type SARS-Cov-2 spike^[17]^ was fused with T4 fibritin (foldon)^[18]^, linked by a flexible GS-linker^[19]^ to generate a trimer structure (**Figure 5A**). For the secretion of this antigen, Secrecon^[20]^, an engineered signaling peptide was used. To evaluate whether this design leads to a soluble, secreting and accurate RBD trimer, DNA fragment coding for N-terminal His-tagged RBD trimer antigen was chemically synthesized and ligated to pcDNA3.1 (+) vector. After protein expression in 293F cells and purification by Histrap immobilized metal affinity chromatography (IMAC), the purified protein was analyzed by SDS-PAGE, in reduced or non-reduced conditions. As illustrated by **Figure 5B**, the non-reduced loading buffer in electrophoresis yielded mainly RBD trimer, while the reduced loading buffer one yielded partly RBD monomer. These results indicated that our RBD-GS linker-T4 fibritin protein design generated RBD trimer definitely. Due to the change of COVID-19 pandemic prevalence from SAR-Cov-2 wild-type to Delta variant, we changed the RBD trimer to Delta variant B.1.617.2, via the mutations of amino acids L452R and T478K in RBD. Next, the Delta RBD trimer was encoded via our cmRNA framework, which contains Anabaena PIE system, Echovirus 29 (E29) IRES, spacer, homology arms, and protein coding region (**Figure 5C**). cmRNA precursor of Delta RBD trimer was *in vitro* transcribed by T7 RNA polymerase, and then cmRNA was generated by circularization reaction (**Figure 5D**) and HPLC-SEC purification (**Figure S4**). The purified cmRNA encoding Delta RBD trimer was transfected to 293 T, and 24 h later cell culturing medium was collected and high level of Delta RBD trimer was detected by ELISA (**Figure 5E**), suggesting that, directed by cmRNA, high expression of soluble and secreting Delta RBD trimer was realized.

**Figure 5.**
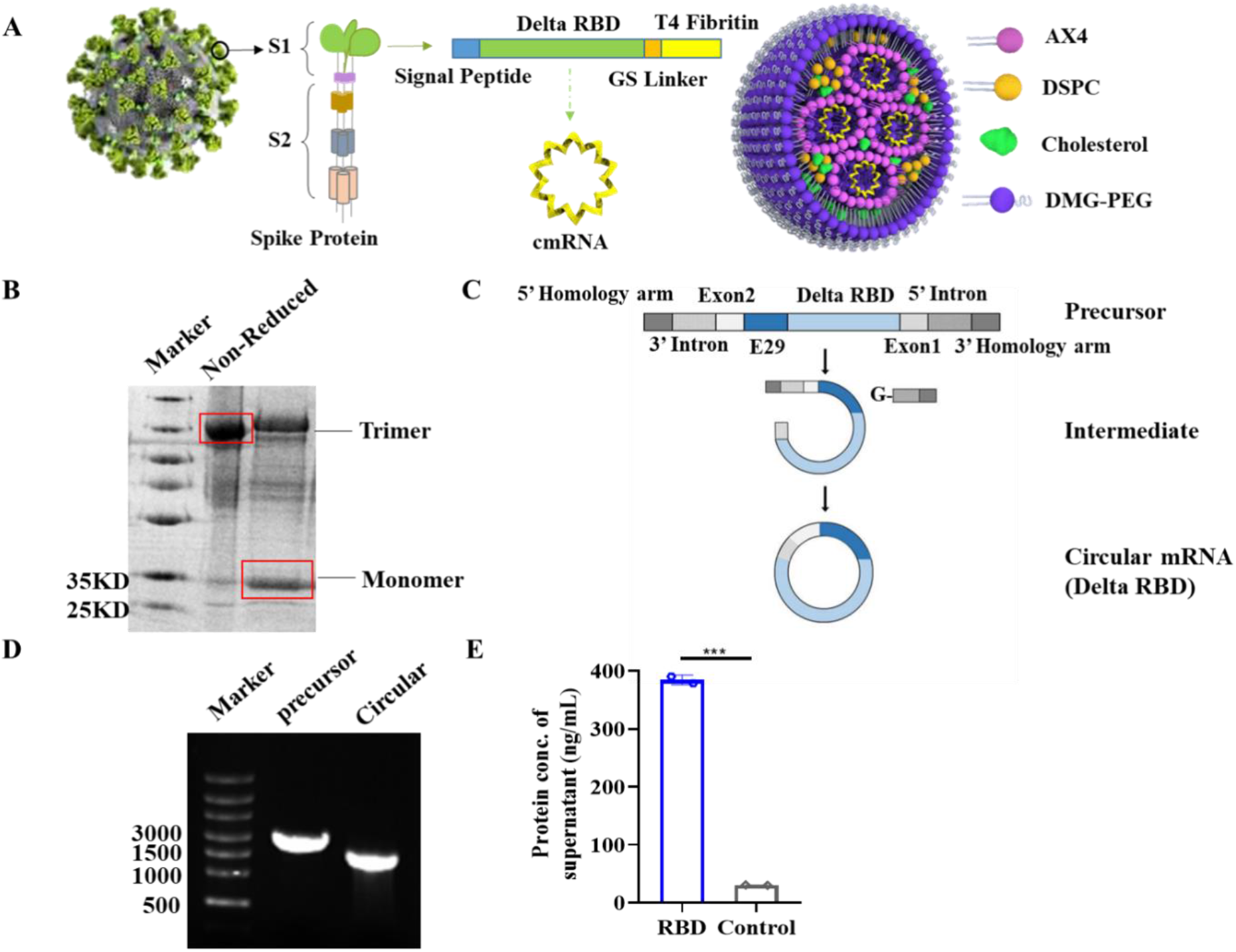
Design and confirmation of cmRNA encoding Delta RBD trimer. (A) Schematic illustration of Delta RBD SARS-Cov-2 mRNA vaccine. (B) RBD trimer and monomer in non-reduced and reduced conditions were analyzed by SAS-PAGE electrophoresis. (C) Schematic illustrtion of cmRNA generation, including the structure of linear RNA percussor, *in vitro* transcription, and circularization. (D) Linear RNA precursor and circular mRNA analyzed by agarose gel electrophoresis. (E) Expression of Delta RBD trimer detected by ELISA. \**P* < 0.05, \*\**P* < 0.01, \*\*\**P* < 0.001, \*\*\*\**P* < 0.0001, n.s. = not statistically significant.

The immunogenicity and efficacy of cmRNA-1130 vaccine that encodes the RBD on spike glycoprotein of B.1.617.2 variant were further evaluated in BALB/c mice. Mice were inoculated via IM injection of cmRNA-1130 containing 30 μg of RBD mRNA, and then boosted with the same dose three weeks later (**Figure 6A**). For the evaluation of safety and efficacy, serums were collected on day 14, 28, 42, 72, 102 and 132, and RBD/spike specific antibodies, SARS-Cov-2 NAbs and RBD specific T cell were measured. No systemic adverse events were observed except for mild swelling at the injection site, which might be a normal reaction to vaccines as a result from temporary innate activation. On day 14 after initial vaccination, mice were phlebotomized for serum NAbs analysis using SARS-CoV-2 surrogate virus neutralization test kit^[21]^, and cmRNA-1130 presented strong serum NAbs with an inhibition value of more than 95%, in sharp contrast with placebo group that induced little immunization (**Figure 6B**). After the second immunization with cmRNA-1130, a high level of NAbs was maintained and the inhibition value was over 95% during the whole experimental period (14-132 d). To further verify the effectiveness of cmRNA-1130, pseudovirus neutralization test (PNA) was used to assess neutralization ability of antibodies generated after second immunization. As indicated by **Figure 6C**, the results showed that mice immunized by cmRNA-1130 demonstrated high Delta pseudovirus neutralization, featured by a notable blockade of the Delta pseudovirus mediated invasion of ACE2-positive cells with a half-maximal inhibitory concentration (IC50) up to 4634 Tu/mL. The results indicated that two doses of cmRNA-1130 vaccination in mice elicited high level and durable anti-SARS-Cov-2 NAbs with strong neutralizing activity against Delta pseudovirus, suggesting that NAbs elicited by RBD trimer are sufficient to blockade full-length spike protein mediated cellular transduction of pseudovirus.

**Figure 6.**
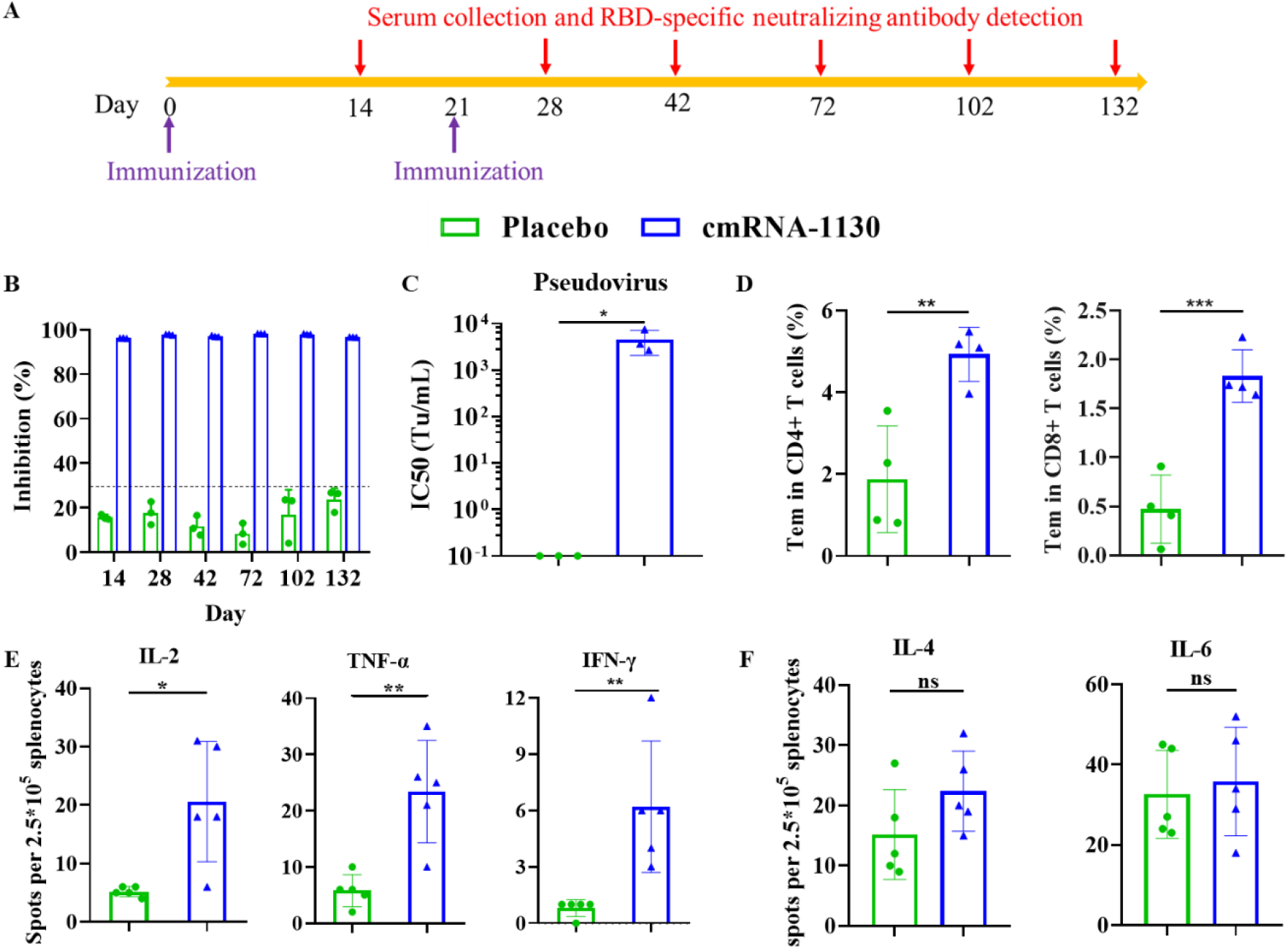
*In vivo* immunization effect of mRNA-AX4-LNP in BALB/c mice. (A) Schematic diagram showing the schedule of immunization and sample collection. (B) The level of RBD-specific neutralizing antibodies in serum of cmRNA-1130 vaccinated BALB/c mice at days 14, 28, 42, 72, 102 and 132. Empty AX4-LNP was used as placebo. (C) Neutralizing antibody (NAb) titers measured using a SARS-CoV-2 B.1.617.1 pseudovirus neutralization assay at day 7 after secondary injection. NAb titers are shown as IC50 values calculated by Reed-Muench method (n = 3). (D) SARS-CoV-2 RBD-specific CD4^+^ and CD8^+^ Tem cells (CD44^+^CD62L^−^) in splenocytes detected by flow cytometry (n = 4). (E) Th1 cytokines including IFN-γ, TNF-α, IL-2 measured by ELISpot assay on day 42 (n = 5). (F) Th2 cytokines including IL-4 and IL-6 in splenocytes measured by ELISpot assay on day 42 (n = 5). Data are shown as mean ± SEM. Significance was calculated using unpaired t test (\**P* < 0.05, \*\**P* < 0.01, \*\*\**P* < 0.001, \*\*\*\**P* < 0.0001, n.s. = not statistically significant.).

Several novel coronavirus vaccines currently under development have been reported to stimulate T cell immune responses in addition to high levels of NAb responses^[22]^. The assessment of cellular immune responses was performed by analyzing T cell activation in the spleen of two doses vaccinated BALB/c mice, on day 21 after the second immunization. **Figure 6D** demonstrated that specific CD4+ cells and CD8+ effector memory T cells (Tem) elicited by cmRNA-1130 were significantly increased compared with placebo group. Cytokines induced by restimulation with the SARS-CoV-2 S1 scanning pool were assessing in splenocytes by ELISPOT assays. Mice treated with cmRNA-1130 revealed much higher expression of T helper 1 (Th1) cytokines tumor necrosis factor α (TNF-α), in which over 20 spot-forming cells per twenty-five thousand splenocytes were detected in the cmRNA-1130 group that were significantly higher than placebo group (**Figure 6E**), as well as other Th1 cytokines including interferon γ (IFN-γ) and interleukin-2 (IL-2). In contrast, no significant difference of T helper 2 (Th2) cytokines including IL-4 or IL-6 cytokines was observed for cmRNA-1130 and placebo groups (**Figure 6F**). The results showed that cmRNA-1130 could activate spike/RBD specific T cells and induce Th1 type specific cell immune response in mice.

The commercialized mRNA vaccines including mRNA-1273 and BNT162b2 are reported to be unstable and easily degradable at room temperature, and must be stored at ultra-low temperature^[23]^. As shown in **Figure S5**, cmRNA-1130 demonstrated little size change, mRNA leakage, and encapsulation efficiency alteration at 4 °C for 6 months as well as 25 °C and 37 °C for 7 days. Besides, cmRNA-1130 could endure repeated freezing and thawing processes, and revealed remarkable stability without obvious change of sizes and mRNA encapsulation after 6 cycles of freezing and thawing. Importantly, cmRNA-1130 following six months’ storage at 4 °C or 6 cycles of freezing and thawing processes still exhibited equivalent NAb level to the fresh one at day 14 post vaccination (**Figure S6A-B**). The superior stability of cmRNA-1130 benefits the efficacy and cold-chain transportation of mRNA vaccines.

## Discussions

We show here a highly potent lipidoid nanoparticle, composed of a leading tail-branched, eight ester bonds included biodegradable lipidoid AX4, is capable of liver-specific delivery of circular mRNA *in vivo*. In comparison with commercialized MC3-LNP, AX4-LNP revealed much faster elimination rate from liver and spleen and induced negligible systemic side effects as characterized by expression of ALT, AST, and CRE. In the previous studies, we developed a new circular mRNA platform, termed cmRNA for efficient and durable expression of various proteins *in vitro and in vivo*^[7c]^. The cmRNA was generated by an RNA polymerase mediated *in vitro* transcription and a step of group I permuted intron-exon (PIE) mediated circularization^[24]^, without biochemical reaction steps such as capping, polyA tail addition and modified nucleoside addition that often required for the manufacturing of linear mRNA. Besides, the cmRNA exhibited longer duration of protein expression, compared to typical linear mRNA, possibly arising from the resistant of cmRNA to exoribonuclease type of RNases^[25]^. Combing the prolonged protein expression by cmRNA and fast elimination rate of AX4-LNP affords mRNA vaccines with outstanding safety and efficacy.

RBD trimer was selected as the vaccine antigen since it has been approved in various pre-clinical studies^[26]^. The T4 fibritin engineered RBD trimer mimics the natural RBD/spike structure of SARS-Cov-2 so as to generate NAbs that target to SARS-Cov-2 spike and avoid the infusion and invasion of this virus into ACE-2 positive cells^[27]^. In the current study, T4 fibritin fusion RBD protein was confirmed as a trimer, and cmRNA encoding Delta RBD trimer was produced and tested *in vitro* to express high level of secreted RBD proteins. Then this cmRNA antigen was encapsulated by AX4-LNP to form a vaccine termed cmRNA-1130. This vaccine elicited high level of Delta SARS-Cov-2 specific NAbs, which were confirmed to neutralize Delta SARS-Cov-2 pseudovirus *in vitro*, and found to maintain for at least 132 days, suggesting a high neutralizing activity and potential long protection duration. According to the previous reports, NAbs that elicited by the current commercial available mRNA vaccines maintain for about 5 months in human^[28]^, comparable to the one elicited by cmRNA-1130 in mice. Considering that the life-span of human is much longer than that of mouse, it is rational to speculate a much longer duration of cmRNA-1130 elicited NAbs in human.

T cell immunity that elicited by mRNA vaccines is one of most advantages over the other kind of vaccines, such as recombinant protein vaccines and inactivated virus vaccines^[29]^. Vaccine derived T cell immunity is based on the expression of antigens in antigen presenting cells (APCs), like dendritic cells, and present T cell epitopes via MHC-I pathway to generate epitope-specific Th1 type CD4^+^ and CD8^+^ T cells^[30]^. In the current study, cmRNA-1130 was found to induced notable RBD/spike specific Th1 type T cell immunity, featured by the identifications of significant up-regulated CD4^+^ and CD8^+^ T cells in splenocytes, as well as the ELISPOT results. The Th1 specific T cell immunity that induced by cmRNA-1130 may be important for the elimination of SARS-Cov-2 virus in human, especially in the conditions of fast evolution of SARS-Cov-2 virus. Mutations of spike protein in SARS-Cov-2 results in the evading from vaccine derived protection, mostly due to the failure or weakness of neutralizing activities of NAbs^[31]^. However, effective T cells, especially memory T cells recognize a broad range of spike/RBD epitopes and battle against viruses directly to avoid the immune evasion of variant viruses, and contribute to remarkable protection against hospitalization or death^[32]^. This explains why mRNA vaccines present higher protection rates against other kinds of vaccines. The successful induction of T cell immunity by cmRNA-1130 suggests that it is a very promising vaccine candidate for the prophylaxis of SARS-Cov-2.

The stability of mRNA vaccine is challenging, due to the intrinsic instability of mRNA molecules. COMIRNATY (BNT162b2), the first FDA approved SARS-Cov-2 mRNA vaccine, has to be stored in an ultra-cold freezer between −90 °C and −60 °C, and the high requirement for its cold-chain transportation may limit its clinical utilization and commercial spaces^[33]^. Therefore, there is a strong demand for the development of high stability mRNA vaccines. In the current study, cmRNA-1130 presented excellent thermal stability, featured by the unaffected vaccine efficacy after storage at 37 °C for 7 days; and good freezing/ thawing stability, characterized by freezing and thawing for 6 times; and good long-term stability, revealed by the storage at 4 °C for 6 months. These results demonstrate that cmRNA-1130 probably can be stored and transported at 4 °C, thus may facilitate its future clinical utilization. The reason for high stability of this vaccine may be the coordination effect of our supreme engineered LNP formulation and the intrinsic high stability of cmRNA. To clarify the exact mechanisms for the high stability of cmRNA-1130, more investigations will be performed in our future studies.

In summary, we have reported on a novel mRNA vaccine (cmRNA-1130) formed from a new biodegradable AX4-LNP and circular mRNA encoding RBD trimer of Delta variant SARS-CoV-2 against SARS-COV-2 Delta variant. cmRNA-1130 possesses some fascinating characteristics: i) AX4 lipid possesses four branched tails and eight ester bonds, facilitating the endosomal escape and mRNA release of mRNA vaccines, and the ultimate elimination of LNP; ii) mRNA-encapsulated AX4-LNP demonstrates liver-targeted protein expression, verifying the potential for the treatment of liver-related diseases; iii) cmRNA provides high stability, efficient and durable expression of various proteins; iv) cmRNA-1130 can elicit potent and sustained neutralizing antibodies against Delta SARS-Cov-2 without causing obvious side effects; v) cmRNA-1130 induces remarkable spike specific Th1-biased T cell responses in mice, resulting in enhanced protection against Delta SARS-Cov-2; vi) cmRNA-1130 vaccine has excellent stability against 6-month storage at 4 °C and freezing-thawing cycles, facilitating the storage and transportation of mRNA vaccines. Thus, mRNA vaccines combing biodegradable AX4-LNP and circular mRNA encoding different proteins emerge as a safe and efficacious protection against various virus variants of COVID-19.

## Materials and Methods

### Synthesis of AX4

N-(3-dimethylaminopropyl)-N′-ethylcarbodiimide hydrochloride (40.9 g, 335.20 mmol) and 4-(dimethylamino) pyridine (8.72 g, 71.47 mmol) were added to the solution of 2-butyloctanoic acid (30.0 g,150.0 mmol) in dichloromethane (400 mL). The reaction was stirred for 5 mins, and 2-propenoic acid 4-hydroxybutyl ester (20.6 g, 143.1 mmol) was added. The mixture was stirred at room temperature for 18 h, and TLC showed a new spot formed. The mixture was diluted with dichloromethane, and washed with a saturated aqueous sodium bicarbonate solution, then brined, dried (MgSO_4_), filtered and evaporated under vacuum. The residue was purified by silica gel chromatography (0-5% ethyl acetate in hexanes) to give 4-(acryloyloxy) butyl 2-butyloctanoate (36.0 g, 100.10 mmol, 77%) as a colorless liquid.

3,3′-Diamino-N-methyldipropylamine (0.74 g, 5.10 mmol) was added to the solution of 4-(acryloyloxy) butyl 2-butyloctanoate (10 g, 30.58 mmol) in methanol (10.0 mL). The mixture was stirred at 40 °C for 18 h. TLC showed a new spot formed, and the mixture was concentrated under vacuum. The residue was purified by silica gel chromatography (0-8% methanol in dichloromethane) to give AX4 (2.0 g, 1.38 mmol, 27 %) as a colorless liquid. ^1^H NMR (400 MHz, CDCl_3_, *δ*): 4.11-4.05 (d, 16H), 2.76-2.72 (d, 8H), 2.43-2.39 (d, 12H), 2.32-2.25 (m, 8H), 2.16 (s, 3H), 1.71-1.17 (m, 84H), 0.87-0.83 (m, 24H).

### Lipid nanoparticle formulation

Lipid nanoparticle (LNP) formulations were prepared through a previously described process. Briefly, the mRNA was diluted in 10 mM citrate buffer (pH 4.0), and the cationic lipid, 1, 2-distearoyl-sn-glycero-3-phosphocholine (DSPC), cholesterol and DMG-PEG were dissolved and mixed in ethanol. The mixture of lipids was then mixed with mRNA solution at the volume ratio of 1:3 using a microfluidic mixer (INano P, Micro & Nano Technology Inc, China). Formulations were then diafiltrated against 10-fold volume of 150 mM PBS (pH7.4) or 20 mM Tris (pH7.4) with 8% sucrose through a tangential-flow filtration (TFF) membrane with 100 kDa molecular weight cut-offs (EMD Millipore), and concentrated to desired concentrations. If needed, then passed through a 0.22-μm filter and stored at 4°C (PBS) or −20°C (20 mM Tris-8% sucrose) until use. All formulations were tested for particle size, distribution, RNA concentration and encapsulation. The concentration and encapsulation rate of mRNA were measured by the Quant-iT™ RiboGreen™ RNA Assay Kit (Invitrogen™ R11490). The size of LNP particles was measured using dynamic light scattering on a Malvern Zetasizer Nano-ZEN 3600 (Malvern).

### Cryo-electron microscopy of mRNA-AX4-LNP

The mRNA-AX4-LNP sample (3 μL) was deposited on a holey carbon grid to form a very thin layer of sample liquid, and was quickly dropped into liquid nitrogen for rapid freezing. After that, the sample was found to transform to a glass state, and finally taken by Cryo-electron microscopy. The Cryo-EM imaging was performed on a Talos F200C equipped with a Ceta 4k x 4k camera, running at an accelerating voltage of 200 kV.

### 6-p-Toluidino-2-naphthalenesulfonic acid (TNS) assay

The pKa of the LNP formulations was determined as previously described. Briefly, a series of buffers with pH ranging from 2.5 to 11 (pH 2.5, pH 3, pH 3.5, pH 4, pH 4.6, pH 5, pH 5.5, pH 5.8, pH 6, pH 6.5, pH 7, pH 7.5, pH 8, pH 8.5, pH 9, pH 9.5, pH 10, pH 10.5, pH 11) were prepared by adjusting the pH of each buffer solution consisting of 10 mM (4-(2-hydroxyethyl)-1 piperazineethanesulfonic acid) HEPES, 10 mM 2-(4-Morpholino) ethane sulfonic acid MES, 10 mM ammonium acetate, 130 mM sodium chloride (NaCl) with 1 N hydrochloric acid (HCl). Additionally, 180 μL of each buffer solution was added to 1.5mL EP tubes, and 6 μL of each LNP formulations (with mRNA) were added to the buffer solutions. Then 4 μL of T NS stock solution (20 μM) in dimethyl sulfoxide (DMSO) was added to the mixture. Next, 180 μL of each above mixture was added to each well of a 96-well plate. The fluorescence intensity was measured at an excitation wavelength of 325 nm and an emission wavelength of 435 nm. The fluorescence intensity was plotted against pH values and fitted using a four-parameter logistic equation (GraphPad Prism). The pH value at which half of the maximum fluorescence is reached was calculated as the pKa of LNP formulations.

### Animal studies

6- to 8-week-old female C57BL/6 and male BALB/c mice were purchased from Vital River (Beijing, China). 6- to 8-week-old female B6/JGpt-ApoEem1Cd82/Gpt mice were purchased form GemPharmatech (Nanjing, China). All protocols were approved by the Guide of Care and Use of Laboratory Animals. Mice were housed in a temperature-controlled, ventilated room and illuminated by artificial light for 12-hour daylight and 12-hour darkness. Animals had free access to food (normal rodent chow) and water.

### Biodistribution of Fluc mRNA-LNP

Male BALB/c mice 5–8 weeks old were purchased from Charles River Laboratories (Beijing, China) and were acclimated for at least 3 days before the initiation of a study. BALB/c mice were IM injected with 10 μg of luciferase-mRNA delivered by LNP. 6 h later, mice were anesthetized with isoflurane gas (2% isoflurane in oxygen, 1 L/min) during injection and imaging procedures. The 150 mg/kg of D-Luciferin (Biosynth AG) was intraperitoneally (IP) injected and mice were imaged in a light-tight chamber using an *in vivo* optical imaging system (IVIS 100; Xenogen Corp.) equipped with a cooled charge-coupled device camera. Bioluminescence images were acquired between 8 and 10 min after D-Luciferin administration. After whole-body imaging, mice were sacrificed and organs were collected for ex vivo imaging. Organs were placed on non-luminescent paper in the imaging chamber and images were acquired as detailed for whole body imaging.

### Lipid clearance in a murine model

Male BALB/c mice were IV injected with 20 μg of mRNA delivered by LNP. 3 mice were sacrificed, the liver and spleen were collected at 2-, 6-, 12-, 24- and 48-h following administration (n=3 mice each time point). The concentrations of AX4 in tissue samples were analyzed by LC-MS/MS and calculated using an established standard curve. Area under curve (AUC) was calculated, which represents accumulative distribution of AX4 in each specimen over 48h-duration post administration.

### pK

BALB/c mice were IV injected with 20 μg of mRNA delivered by LNP. Then, about 50 μL of blood samples were taken from the mouse orbital at indicated time points (0.083, 0.25, 0.5, 1, 2, 4, 8, 12, 24 and 48 h) and centrifuged at 3500 rpm for 5 min. The supernatant was added to 0.3 mL of ethanol and AX4 was extracted at 4 °C overnight, followed by centrifugation for 10 min at 5000 rpm. Finally, the content of AX4 in the supernatant was measured by LC-MS/MS. The elimination half-life (t_1/2_, β) was calculated by fitting the experimental data using software Origin 9 exponential decay 2 model: y = A_1_ × exp(−x/t_1_) + A_2_ × exp(−x/t_2_) + y_0_, and then taking t_1/2_, β = 0.693 × t_2_.

### Quantification of lipid by LC-MS/MS

Blood and tissue samples were homogenized by vortex following the addition of ethanol. Lipid was extracted and analyzed against calibration standards prepared in a matching blank. Chromatographic separation and quantification were accomplished with a liquid LC-MS/MS system. Samples were separated on a ACQUITY UPLC BEH C18 column (1.7 μm, 2.1 × 100mm; Waters Corporation) equilibrated with moving phase (H_2_O: MeOH: FA, 50:50:1) (Merck Corporation). A triple-quadrupole MS/MS system (AB Sciex, QTRAP 5500) operated in positive ion mode was used for signal detection.

### Gene cloning and vector construction

DNA fragments that contain PIE elements, IRES, coding regions and others were chemically synthesized and cloned into pUC57 plasmid vector. For RBD trimer protein expression, DNA chemically was synthesized and cloned into the vector pcDNA3.1 (+). DNA synthesis and gene cloning were customized ordered from Jiangsu Genscript Co.Ltd. (Nanjing, China).

### cmRNA preparations

cmRNA precursors were synthesized by *in vitro* transcription from a linearized plasmid DNA template using a Purescribe™ T7 High Yield RNA Synthesis Kit (CureMed, Suzhou, China), followed by DNase I (CureMed, Suzhou, China) treatment for 15 min to remove DNA template. Afterward, unmodified linear mRNA was column purified using a GeneJET RNA Purification Kit (Thermo Fisher). For circularization, GTP was added to a final concentration of 2 mM along with a buffer including magnesium (50 mM Tris-HCl, (pH 8.0), 10 mM MgCl_2_, 1 mM DTT; Thermo Fisher). RNA was then heated to 55 °C for 15 min, and then column purified. For high-performance liquid chromatography (HPLC), RNA was run through a 30 × 300 mm size exclusion column with the particle size of 5 μm and pore size of 1000 Å (Sepax Technologies, Suzhou, China) on an SCG protein purification system (Sepure instruments, Suzhou, China). RNA was run in RNase-free Phosphate buffer (pH:6) at a flow rate of 15 mL/minute. RNA was detected and collected by UV absorbance at 260 nm. Concentrate the purified cmRNA in an ultrafiltration tube, and then replace the phosphate buffer with an RNase-free water.

### Cell lines

Human cell line HEK293T were cultured in DMEM (BI) supplemented with 10% fetal calf serum (Gibco) and penicillin/streptomycin antibiotics (100 U/mL penicillin, 100 μg/mL streptomycin, Gibco). HEK293F cells were maintained in Free Style™ 293 Expression Medium. All cell lines were maintained at 37°C, 5% CO_2_, and 90% relative humidity.

### mRNA *in vitro* transfection

Cells reaching 60%~80% confluent were transfected with mRNA using Lipofactamine MessengerMAX (Invitrogen) according to the manufacture’s instruction. Briefly, the mRNA and Lipofactamine Messengermax reagent were diluted with Opti-MEM (Gibco), mixed together and incubated for 5 min at room temperature for complex formation. Then the entire mixture was added to each well and the cells were incubated in the in a 5% CO2 incubator.

### Protein expression and purification

The plasmid pcDNA3.1(+)-RBD for the expression of RBD trimer was transfected to HEK293F by 25 kd PEI (Shanghai Maokang Co.,Ltd).900 μg of plasmid was mixed with 2.7 mL PEI (1mg/mL) and 45 mL Opti-MEM medium and added to 150 mL 293F cells, with cell density as 1.5 × 10^6^/mL. After 3 h incubation at 37 °C, 150 mL Free Style™ 293 Expression Medium was supplemented, and cells were incubated for 4 days in the condition of 37 °C, 220 rpm, 5% CO_2_. Cell cultured medium was collected by 8000 rpm, 20 min centrifugation, and protein was purified by Histrap FF column (Cytiva) in immobilized metal affinity chromatography (IMAC). The purified protein was analyzed by 10% SDS-PAGE with reduced or non-reduced loading buffer.

### Biochemical evaluation of liver and kidney function

For liver toxicity studies, whole blood samples were collected from the posterior orbit to clean dry centrifuge tubes and centrifuged at 5000 rpm for 15 min, and serum was separated and collected, then kept at −80 °C for further biochemical analyses of alanine transaminase (ALT), aspartate transaminase (AST) and creatinine (CRE) using the commercially available kits (Laboratory Biodiagnostics Co., Cairo, Egypt). All procedures were performed according to the manufacturer’s instructions.

### Enzyme-linked immunosorbent assay

Cellular immune responses in the vaccinated mice were assessed using IFN-γ, TNF-α, IL-2, IL-4, or IL-6 precoated ELISPOT kits (MabTech), according to the manufacturer’s protocol. Briefly, the plates were blocked using RPMI 1640 (Thermo Fisher Scientific) containing 10% FBS and incubated for 30 minutes. Immunized mouse splenocytes were then plated at 500,000 cells/well, with SARS-CoV-2 S1 scanning pool (2 μg/ml of each peptide in the cell culture). After incubation at 37 °C, 5% CO_2_ for 48 h, plates were washed with wash buffer and biotinylated anti-mouse IFN-γ, TNF-α, IL-2, IL-4 or IL-6 antibody was added to each well followed by incubation for 2 h at room temperature. Subsequently, each well was wash and incubated with streptavidin-HRP for 1 h at room temperature, following by the addition of TMB substrate solution. The air-dried plates were read using the automated ELISPOT reader AID ELISPOT (AID) and the numbers of spot-forming cells (SFC) per 500,000 cells were calculated.

### Flow cytometry analysis

T cell proliferation in immunized mice were evaluated using a FACS Calibur flow cytometer (BD Biosciences). Briefly, a total of 1,000,000 mouse splenocytes were stimulated with SARS-CoV-2 S1 scanning pool (2 μg/ml of each peptide) for 2 h at 37 °C with 5% CO_2_. Splenocytes were twice washed with PBS and stained with fluorescently conjugated antibodies to CD3 (APC) (BioLegend), CD4 (Brilliant Violet 605™) (BioLegend), CD8 (Brilliant Violet 421™) (BioLegend), CD44 (PE) (BioLegend) or CD62L (FITC) (BioLegend). Dead cells were stained with eBioscience™ Fixable Viability Dye eFluor™ 780 (Thermofisher). Data are analyzed with FlowJo software.

### SARS-CoV-2 Surrogate virus neutralization assay

The neutralizing activity of mouse serum samples was detected by SARS-CoV-2 Surrogate Virus Neutralization Test Kit (L00871, GenScript). For the commercial kit, the assay was performed according to the manufacturer’s instructions. In short, serum samples were diluted with dilution buffer and preincubated with HRP-RBD in a 1:1 ratio for 30 min at 37°C and then added to the capture plate in wells precoated with hACE2. After 15 min of incubation at 37 °C, plate was washed four times with wash buffer. TMB solution was added and incubated for 15 min at room temperature in the dark. After incubation, stop solution was added to mixture and promptly analyzed. Optical density at 450 nm (OD450) was measured using a SpectraMax Paradigm microplate reader (Molecular devices).

### *In vivo* biodistribution and luciferase mRNA delivery studies

Male BABL/c mice received injections of LNP containing 10 μg Fluc mRNA by IMinjection and luciferase imaging was conducted on an IVIS (PerkinElmer, Waltham, MA). Female WT C57BL/6 mice or B6/JGpt-ApoEem1Cd82/Gpt (ApoE^−/−^) mice received injections of LNP by IV injection. 10 min before imaging, mice were IP injected with D-luciferin (150 mg/kg) and anesthetized with isoflurane in oxygen. For co-localization studies, mice were injected with LNP containing Fluc mRNA labeled with Cy5 by IM injection. Before imaging, mice were performed as previous described, and then imaged for both bioluminescence and Cy5 fluorescence at an excitation/emission = 649/670 nm. Next, mice were sacrificed and dissected, and the liver, kidneys, heart, lung, and spleen were imaged for bioluminescence and Cy5 fluorescence. Image analysis was conducted using Living Image software.

### Stability of cmRNA-1130

For the evaluation of long-term stability, cmRNA-1130 was incubated at 4, −20 or −80°C for 6 months. In addition, male BALB/c mice aged 6-8 weeks were inoculated with 30 μg of the incubated mRNA-LNP via IM route. At day 14 after immunization, mice were phlebotomized for serum neutralizing antibody analysis using SARS-CoV-2 Delta variant surrogate virus neutralization test kit (Genscript). For the evaluation of freeze-thaw stability, cmRNA-1130 was incubated at −20 or −80°Cand underwent 6 times freezing and thawing. For the evaluation of thermal-stability, mRNA-LNP was incubated at 4, 25 or 37°C for 1, 4, and 7 days. All formulations of cmRNA-1130 were tested for size, PDI, concentration and EE.

## Supporting information

**Figure S1.**
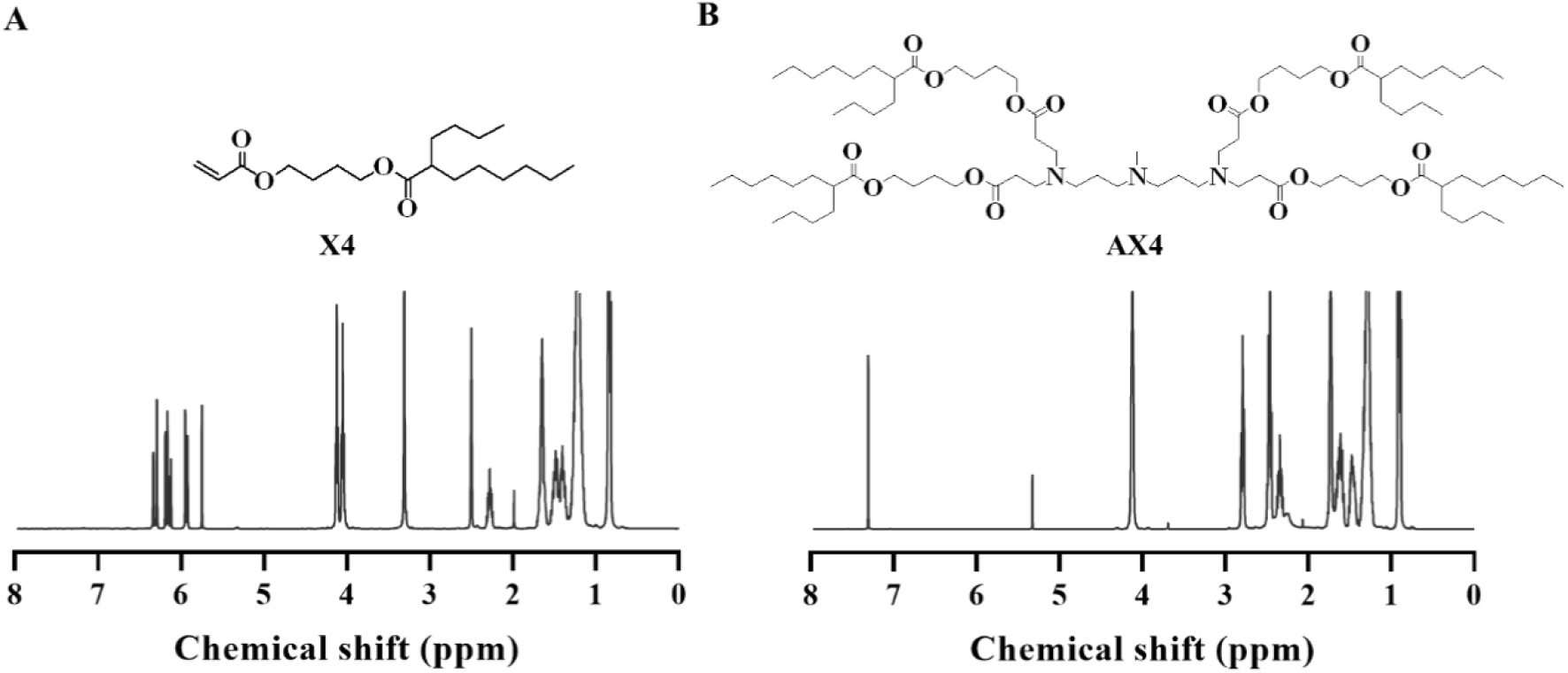
^1^H NMR spectra of X4 (400 MHz, DMSO-*d*_*6*_) (A) and AX4 (400 MHz, CDCl_3_) (B).

**Figure S2.**
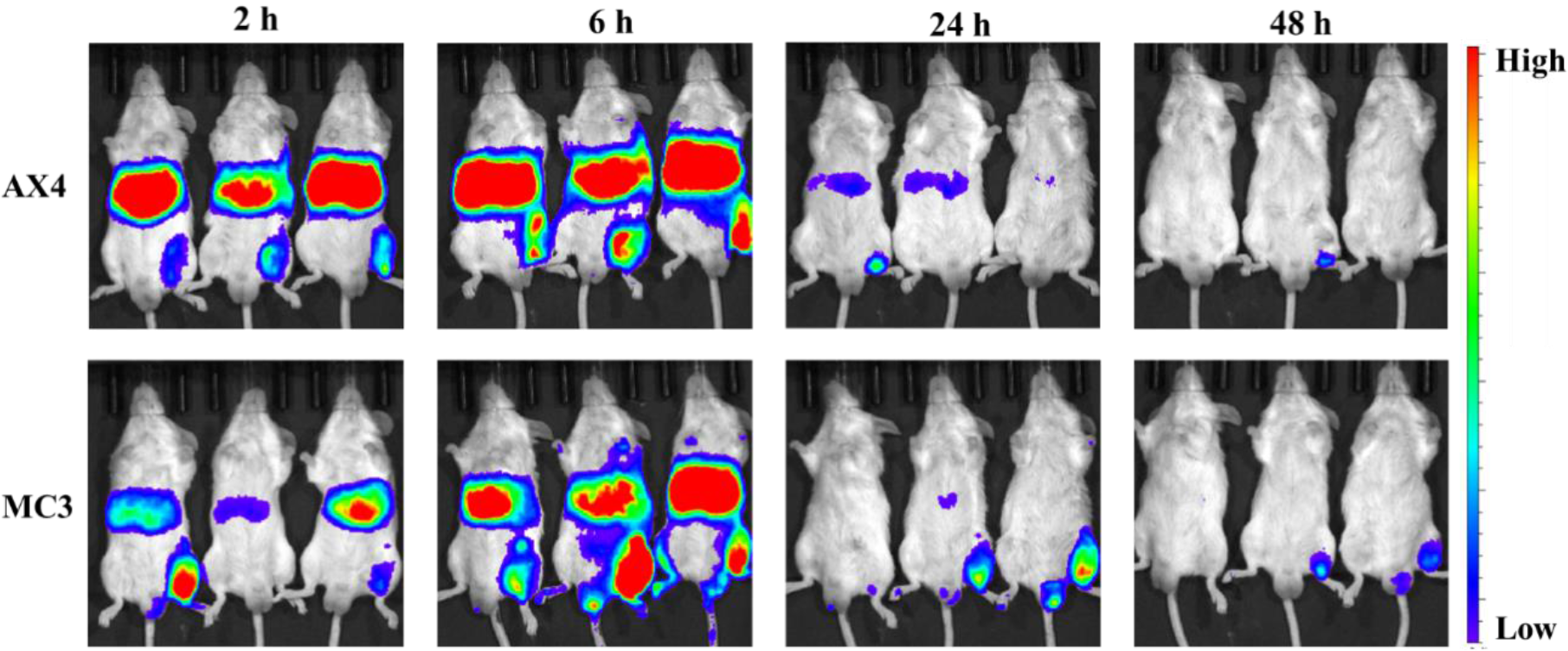
*In vivo* imaging of BALB/c mice following IM administration with Fluc mRNA-encapsulated AX4 and MC3 LNP at a dose of 0.5 mg mRNA equiv./kg.

**Figure S3.**
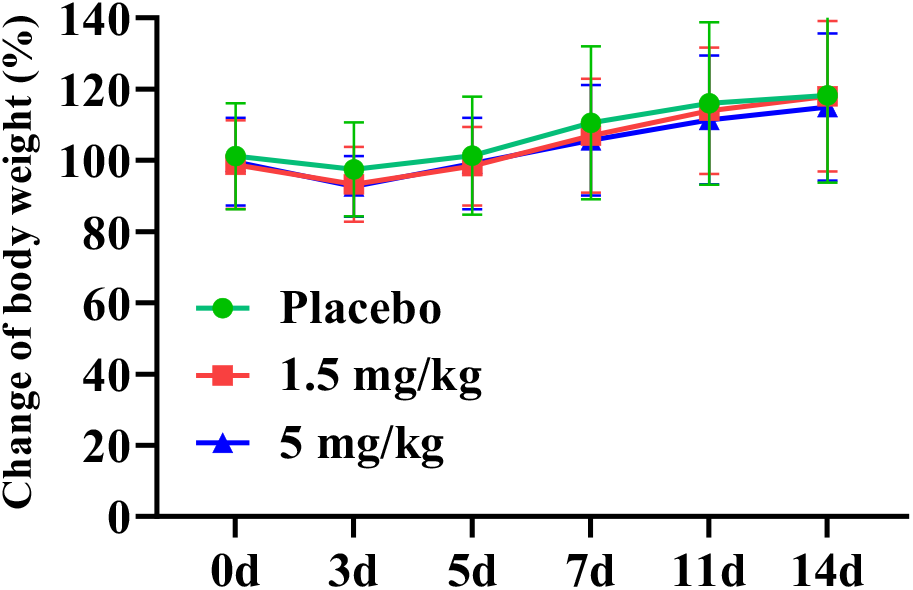
Change of body weight of mice after IM administration of different doses of mRNA-LNP.

**Figure S4.**
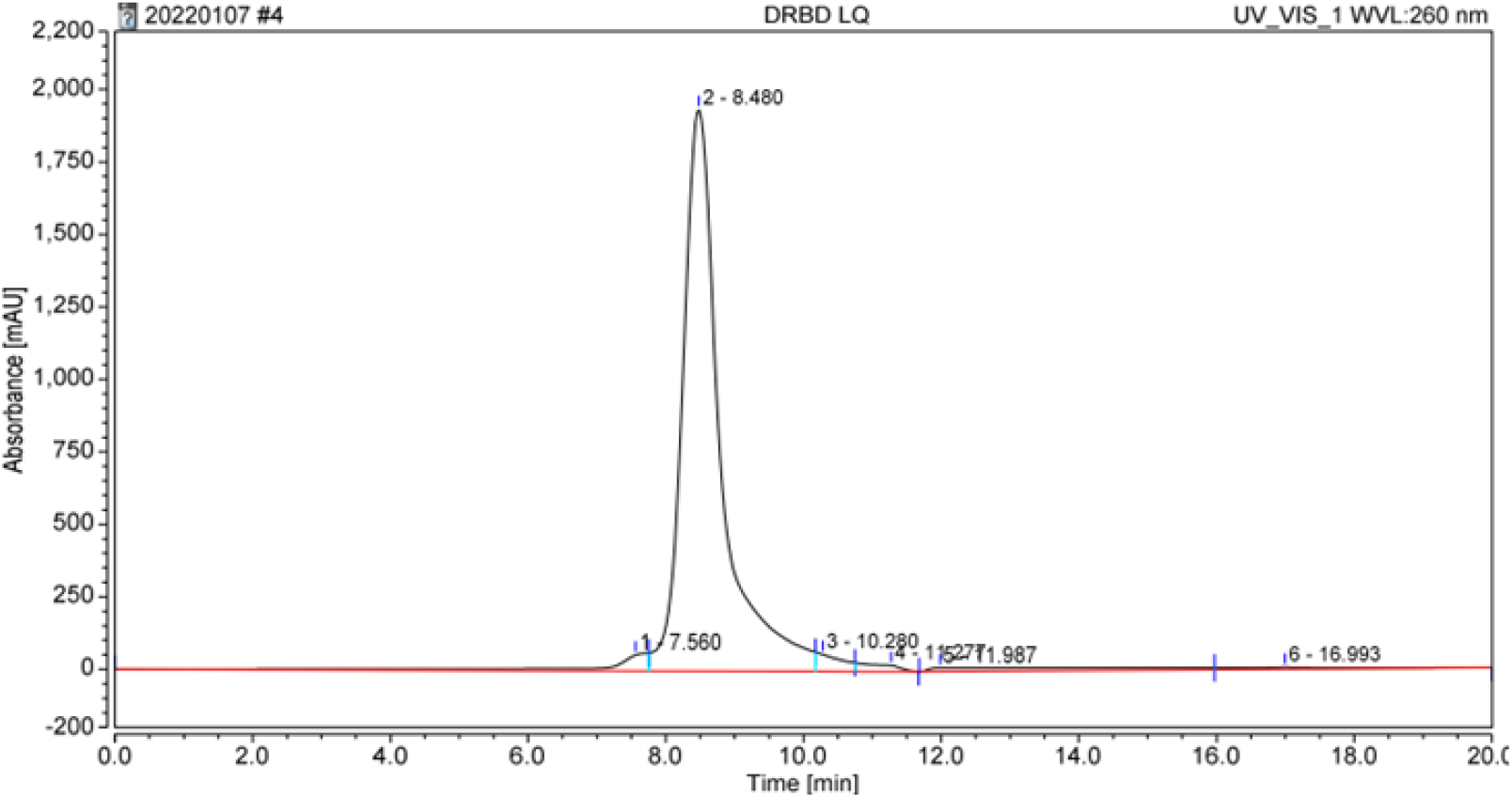
Analysis of the purity of cmRNA that encodes Delta RBD trimer by HPLC-SEC.

**Figure S5.**
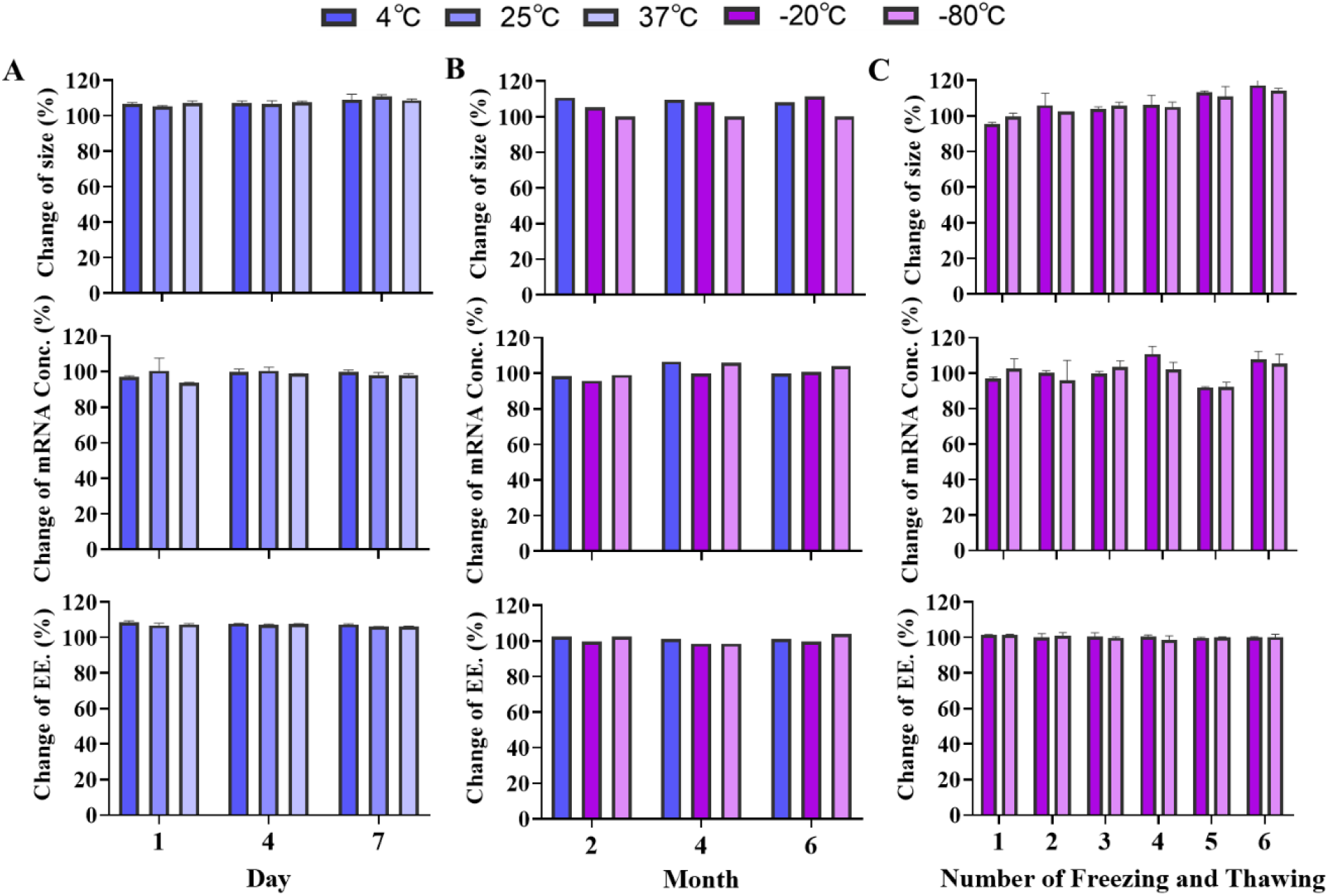
Stability characterization of cmRNA-1130. (A) Changes of size, mRNA concentration, and EE. of cmRNA-1130 following incubation at 4 °C, 25 °C, 37 °C for 1, 4, 7 days. (B) Changes of size, mRNA concentration, and EE. of cmRNA-1130 following incubation at 4 °C, −20 °C, −80 °C for 2, 4, 6 months. (C) Changes of size, mRNA concentration, and EE. of cmRNA-1130 following 6 times of freezing and thawing cycles at −20 °C and −80 °C.

**Figure S6.**
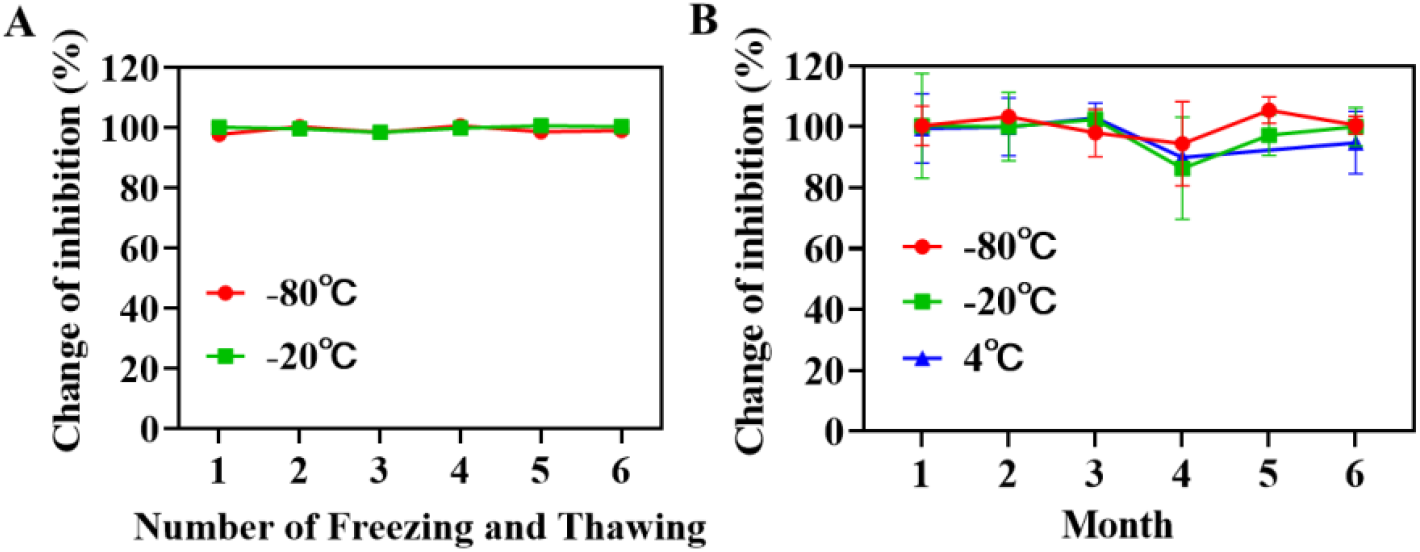
Changes of neutralizing antibodies generated by cmRNA-1130 following (A) 6 times of freezing and thawing cycles at −20 °C and −80 °C, and (C) the storing at 4 °C, −20 °C and −80 °C for 1, 2, 3, 4, 5, 6 months. Serum level of RBD-specific neutralizing antibodies in BALB/c mice measured at 14 day-post single administration of cmRNA-1130.

